# Anti-CD137 monoclonal antibody enhances trastuzumab-induced, natural killer cell-mediated cytotoxicity against pancreatic cancer cell lines with low human epidermal growth factor-like receptor 2 expression

**DOI:** 10.1101/361279

**Authors:** Takushi Masu, Masanori Atsukawa, Katsuhisa Nakatsuka, Masumi Shimizu, Daishu Miura, Taeang Arai, Hirotomo Harimoto, Chisa Kondo, Keiko Kaneko, Seiji Futagami, Chiaki Kawamoto, Hidemi Takahashi, Katsuhiko Iwakiri

## Abstract

**Background:** Because human epidermal growth factor-like receptor (HER) 2 is expressed on the surface of human pancreatic carcinoma cells to varying degrees, trastuzumab, an anti-HER2 monoclonal antibody (mAb), is expected to exert antibody-dependent, natural killer (NK) cell-mediated cytotoxicity (ADCC) against the cells. However, some reports found that the effect of trastuzumab against human pancreatic carcinoma cells was limited because most express only limited HER2.

**Methods:** We examined whether anti-CD137 stimulating mAb could enhance trastuzumab-mediated ADCC against Panc-1, a human pancreatic cancer cell line with low HER2 expression, in vitro.

**Results:** Supplementation of anti-CD137 mAb could improve trastuzumab-mediated ADCC against Panc-1 which was insufficient without this stimulating antibody. The ADCC differed in individual cells, and this was related to the expression of CD137 on the surface of NK cells after trastuzumab stimulation in association with the Fcγ-RIIIA polymorphism. NK cells with Fcγ-RIIIA-VV/VF showed high levels of ADCC against Panc-1, but those with Fcγ-RIIIA-FF did not show optimal ADCC. In addition, trastuzumab-mediated ADCC against the human pancreatic cancer cell line Capan-1 with high HER2 expression was generally high and not affected by the Fcγ-RIIIA polymorphism.

**Conclusions:** These results demonstrated that in Fcγ-RIIIA-VV/VF-carrying hosts, trastuzumab plus αCD137 mAb could induce effective ADCC against HER2-low-expressing pancreatic cancer cells. This also indicates the therapeutic potential for unresectable human pancreatic cancer, in which HER-2 expression is generally low.

## Introduction

Pancreatic carcinoma is difficult to cure [1], and the prognosis of unresectable pancreatic cancer patients is very poor [2]. Although various attempts have been made to establish innovative therapeutic regimens, the efficacy of current chemotherapy regimens remains inadequate [3–8]. Among the chemotherapy regimens used to treat unresectable pancreatic carcinoma, gemcitabine-based ones are common because they maintain the quality of the remaining life of patients without serious complications. Among newly established regimens, the combination of gemcitabine plus aluminum-bound (nab)-paclitaxel was reported to increase the mean survival interval (MSI) from 6 to 10 months compared with gemcitabine alone [7]. Furthermore, the FOLFILINOX regimen greatly improves the MSI of patients with unresectable pancreatic carcinoma, although many patients fail to complete this regimen because of its serious side effects [8]. Thus, the clinical efficacy of these regimens should be improved and new strategies for the treatment of pancreatic carcinoma are needed.

Trastuzumab (Tmab) is a specific monoclonal antibody (mAb) against human epidermal growth factor-like receptor (HER) 2 [9] expressed on various tumor cells [1–14], especially in breast [10] and gastric carcinoma [11]. Antigen-dependent cell-mediated cytotoxicity (ADCC) is the initial mechanism of action of Tmab [15, 16], and there are many reports on the clinical efficacy of Tmab against HER2-expressing tumors, especially against breast carcinoma [17–21]. HER2 is also expressed in varying levels on the surface of human pancreatic carcinoma cells [22, 23], and some reports indicated that Tmab induces ADCC against human pancreatic cancer in vitro and in vivo [24–28]. However, the clinical efficacy of Tmab against human pancreatic carcinoma is inadequate [24] because it was usually investigated in HER2-high-expressing cell lines [26–28], whereas most human pancreatic cancers express only low levels of HER2 [22]. Hence, the clinical efficacy of Tmab against human pancreatic carcinoma remains controversial.

Recently, some groups have tried to up-regulate Tmab-mediated ADCC with the addition of various monoclonal antibodies [29–31]. Notably, Kohrt HE et al. [32] and Houot R et al. [33] reported that anti-CD137 mAb (αCD137) could enhance the Tmab-mediated ADCC against human breast cancer cells. CD137 (4-1BB) is known to act as a co-stimulatory molecule in combination with Fc receptor-mediated stimulatory signaling [34] and is expressed on the surface of natural killer (NK) cells after stimulation [35]. Thus, the hypothesis that the addition of αCD137 to Tmab could up-regulate ADCC against HER2-low-expressing target cells was put forward.

Based on that hypotheses and previous findings, we investigated the effects of αCD137 for NK cell activation to up-regulate Tmab-mediated ADCC against HER2-low-expressing human pancreatic carcinoma cell lines as part of efforts to
a new regimen for unresectable human pancreatic carcinoma.

## Materials and Methods

### Cell lines and cultures

Human pancreatic carcinoma cell lines Panc-1 (HER2-low-expressing cell line), Capan-1 (HER2-high-expressing cell line), and the NK cell-sensitive thymoma cell line K562 were purchased from the American Type Culture Collection (Manassas, VA). HER-2 expression on the surface of Panc-1 and Capan-1 was confirmed by flow cytometry (Figure 1-A). All these cell lines were maintained according to the manufacturer’s instructions. In brief, Panc-1 was cultured with Dulbecco’s modified Eagle’s medium (DMEM, Gibco Life Technologies, Santa Clara, CA) supplemented with 10% heat-inactivated fetal calf serum (FCS) and 10% penicillin and streptomycin solutions (Gibco). Capan-1 was cultured with ISCOVE modified Eagle’s medium (Sigma Chemical Company, St. Louis, MO) supplemented with 10% FCS. Cells were removed by short-term incubation with trypsin-EDTA (Gibco) and about one-third to one-fourth of the viable cells were removed and resuspended in fresh culture medium twice weekly. K562 was cultured in DMEM supplemented with 10% heat-inactivated FCS, HEPES-buffer solution 5 mM, penicillin and streptomycin solutions 100 U/ml, L-glutamine 2 mM, sodium pyruvate solution 2 mM, and nonessential amino acid solution 2 mM (all purchased from Gibco-BRL), modified vitamins 2 mM (DS Pharma, Osaka, Japan), and 2-mercaptoethanol 2 mM (Sigma Chemical).

**Figure 1.**
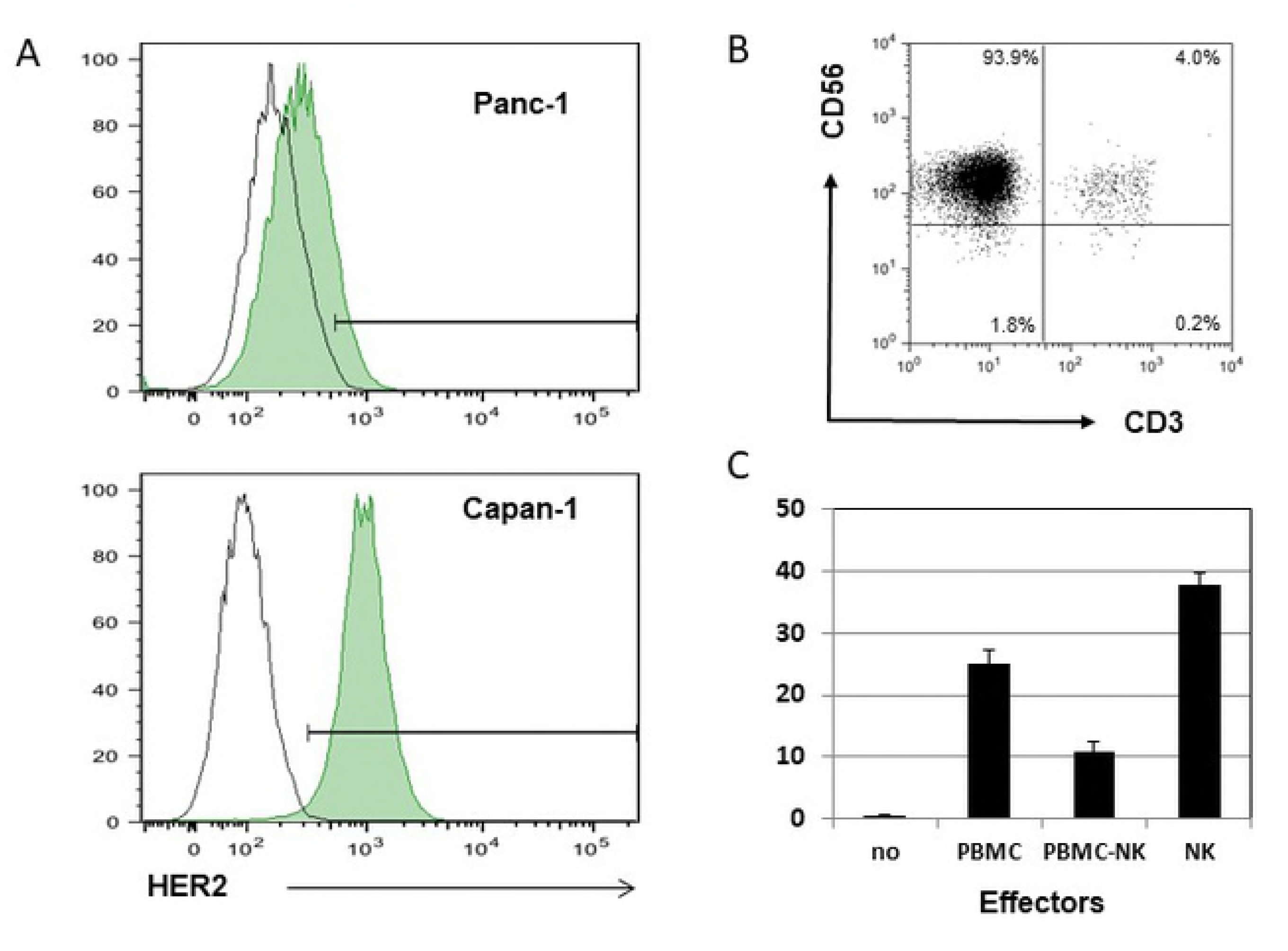
A: Flow cytometric analysis was performed to confirm HER2 expression on Panc-1 (upper panel) and Capan-1 (lower panel). B: The isolated NKs cell were confirmed to be CD3 negative and CD56 positive by flow cytometry. C: The fundamental viability of the isolated NK cells was evaluated as cytotoxicity against the NK-sensitive thymoma cell line K562.

### Monoclonal antibodies

Tmab was kindly provided by Roche-Chugai Japan Co. Ltd. (Tokyo, Japan). It was suspended in phosphate-buffered saline (PBS) 100 μg/ml and stored at –80°C until use. Goat-poly anti-human CD137 stimulation Ab (αCD137, R&D Systems, Minneapolis, MN) was also suspended in PBS 10 μg/ml and stored at −80°C until use.

### Preparation of human NK cells

Peripheral blood was obtained from 12 healthy individuals who were serologically confirmed to be free from hepatitis B and C and human immunodeficiency virus infection. This study protocol conformed to the ethical guidelines of the Declaration of Helsinki as reflected in a priori approval by the Institutional Review Committee of Nippon Medical School. Human NK cells were purified from peripheral blood mononuclear cells (PBMCs) isolated from heparinized blood using the Ficoll Paque (Amersham, Buckinghamshire, UK) density-gradient method. The EasySep human NK cell enrichment kit (Stemcell Technologies, Vancouver, Canada) was used to separate human NK cells according to the manufacturer’s instructions. Briefly, PBMCs were suspended in the recommended buffer and incubated with tetrameric antibody complexes recognizing CD3, CD4, CD14, CD19, CD20, CD36, CD66b, CD123, HLA-DR, and glycophorin A for 10 min at 4°C. Dextran-coated magnetic particles were then added, and the mixture was incubated for an additional 5 min at 4°C. All fractions including the coated cells were placed into the EasySep magnetic chamber for 2.5 min, and those unlabeled were collected as the NK cell-enriched fractions. Cells were washed with PBS and resuspended in DMEM supplemented with 10% FCS and 10% penicillin and streptomycin solutions (assay medium). The purity of NK cells collected in this process was greater than 90% as confirmed by the expression of CD3 and CD56 in flow cytometry (Figure 1-B). The cytotoxic activity of each NK cell was titrated against that of K562 (Figure 1-C). NK cells were resuspended in culture media and used for the following assays.

### Cell culture procedures

1 × 10^6^/well of each target cell line was suspended in 1 ml of assay medium and sheeted in a 48-well flat-bottom culture plate. Tmab 0 to 10 μg/ml was added to each well and incubated for 0, 2, and 12 h at 37°C. Cells were harvested and resuspended in assay medium, and 5 × 10^3^/well of each target cell line was sheeted in a 96-cell U-bottom plate and incubated with 10-to 40-fold amounts of NK cells for 4 h at 37°C. Anti-CD137 mAb 10 μg/ml was added to some wells to examine whether it could enhance Tmab-induced ADCC. After incubation, the plates were centrifuged for 1 min at 1000 rpm, and culture supernatants were collected for cytotoxicity assays and cytokine titration assays. The cells were harvested and used for evaluation of the kinetics of cell surface molecules.

### Flow cytometry

Flow cytometric analysis was performed using a FACSCant-II (BD-Bioscience, San Jose, CA). For staining cell surface molecules, 500,000 cells were harvested, washed twice with PBS, and pelleted. The following antibodies were used: fluorescein-isothiocyanate (FITC)-conjugated anti-human CD3; phycoerythrin (PE)-conjugated anti-human CD56 (BD Bioscience); PE-conjugated anti-human CD16 (clone 3G8), CD137, PD1, and NKG2D (Biolegend, San Diego, CA); PE-conjugated anti-human CD16 (clone MEM154, Immunological Sciences, Rome); and FITC or PE-conjugated mouse IgG1, Isotypecontrol (Biolegend). Propidium iodide was used to confirm the percentage of dead cells. One hundred thousand events were acquired for each sample and analyzed using FlowJo software (Tree Star Inc., Ashland, OR).

### Determination of FcRγIII polymorphism

The FcRγIIIA polymorphism of NK cells was evaluated according to the previous reports [36, 37]. In brief, freshly isolated NK cells were harvested and stained separately with two anti-FcRγIIIA mAbs: clone 3G8 and clone MEM-154. The 3G8 mAb binds to a nonpolymorphic epitope of FcRγIIIA, whereas binding of MEM-154 mAb is dependent on the valine expression of FcRγIIIA. The percentage of cells stained positively for each Ab was counted, and the ratio of MEM-154 positive cells/3G8 positive cells was calculated. The FcRγIIIA polymorphism was determined using the formula V/V: MEM154/3G8 >0.62, F/F: <0.04, and F/V: ratio between 0.15 and 0.48.

### Cytotoxicity assay

The CytoTox 96 nonradioactive cytotoxicity assay kit (Promega, Madison, WI) was used for evaluating NK cell cytotoxicity according to the manufacturer’s instructions. In brief, 5 × 10^3^ treated target cells were resuspended in 50 μl of assay medium and placed in a 96-well round-bottom culture plate. Ten- to 40-fold amounts of NK cells suspended in 50 μl of assay medium were added to each well and also sheeted in empty wells to measure the spontaneous lysis of effector cells. Lysis solution 50 μl and assay medium 50 μl were sheeted in another well for measurement of maximal and minimal lysis. The plate was centrifuged at 250 g for 4 min and incubated for 4 h at 37°C. Lysis solution 10 μl was added to each well 45 min before the end of incubation. The plate was centrifuged again for 4 min at 250 g, and then culture supernatants were collected and replaced in a 96-well flat-bottom assay plate. Substrate mixed buffer 50 μl was added to each well and incubated for 30 min at room temperature while shaded from light. Stop solutions were added to each well at the end of incubation. The amount of lactate dehydrogenase (LDH) released in the supernatant was measured as absorbance at 490 nm using an ELISA reader (Bio-Rad, Hercules, CA). The percentage of specific cytotoxicity was calculated by the following formula: Percent cytotoxicity = (Sample-Effector-minimal)/(Maximal-minimal) × 100.

### Measurement of cytokines

NK cells were plated at 1 × 10^6^/ml in a 48-well plate and stimulated with and without Tmab 10 μg/ml for 48 h at 37°C. Culture supernatants were collected and stored immediately at –80°C. Enzyme-linked immunosorbent assays (ELISA) were performed to titrate interferon (IFN)-γ and tumor-necrosis factor (TNF)-α in the culture supernatants using DUOSET anti-human IFN-γ and TNF-α ELISA kits (R&D Systems).

### Statistics

The paired t-test and Man-Whitney U-test were performed to determine the significance of differences between groups in this study using GRAPHPAD PRISM (GraphPad Software, La Jolla, CA). All experiments were repeated five times, and a p value of < 0.05 was considered to represent a statistically significant difference.

## Results

### Tmab-mediated ADCC against the HER2-high-expressing human pancreatic cancer cell line

First, the Tmab-mediated ADCC against Capan-1, the HER2-high-expressing human pancreatic cancer cell line, was confirmed. As shown in Figure 2-A, while NK cells, isotype IgG, and Tmab did not demonstrate cytotoxicity against Capan-1, 10 μg/ml of Tmab added to NK cells lysed Capan-1. To determine the optimal conditions for inducing Tmab-mediated ADCC, the indicated concentrations of Tmab were examined, and no difference was seen between any of them (Figure 2-B). The necessity for Tmab pretreatment of Capan-1 was also examined, and the results indicated that long-term pretreatment with Tmab decreased the ADCC against Capan-1 (Figure 2-C). Based on those results, we performed the following ADCC assay with 10 μg/ml of Tmab without pretreatment of target cells.

**Figure 2.**
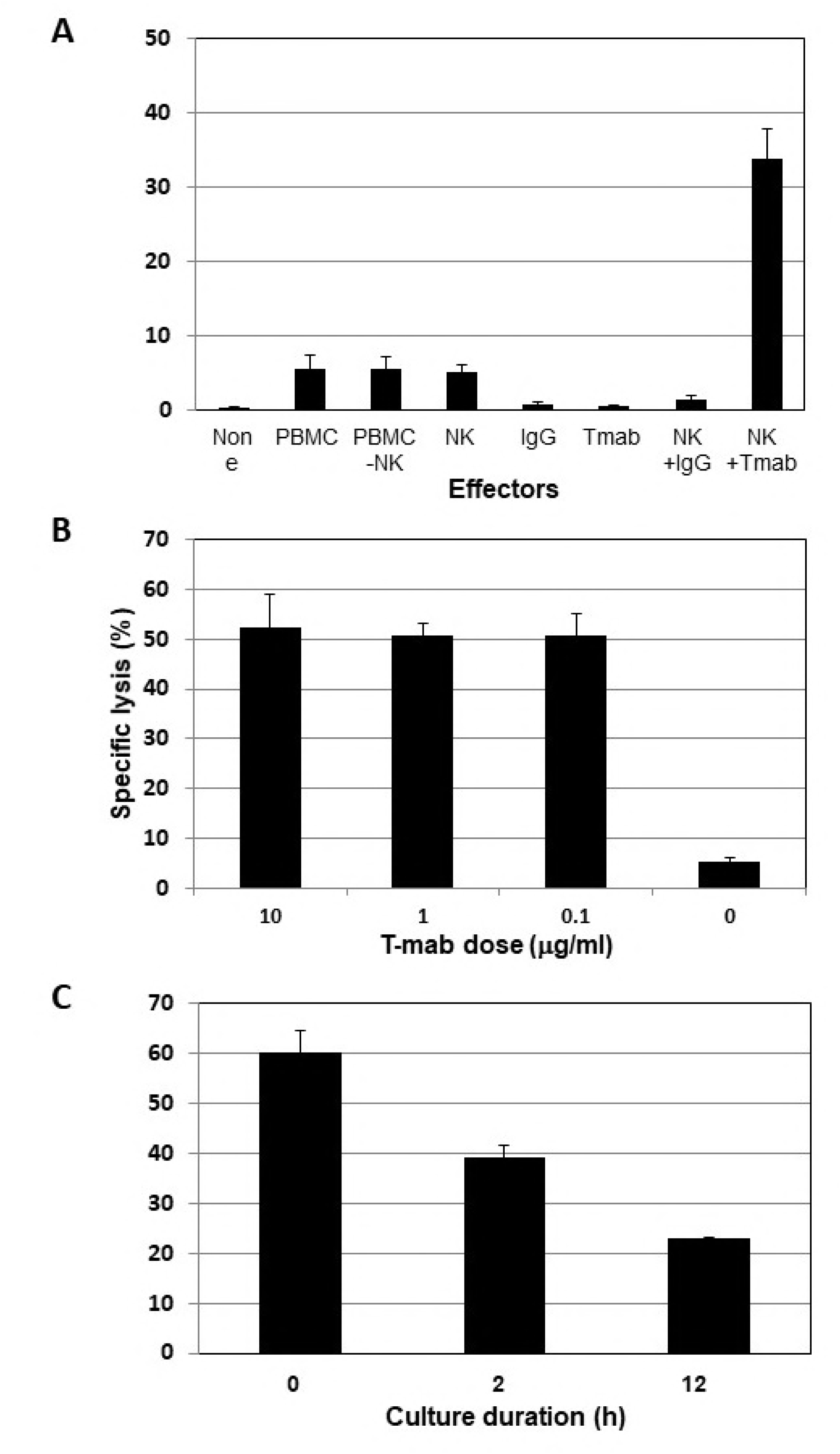
LDH-releasing assays were performed to confirm the appropriate conditions for Tmab-mediated ADCC against Capan-1, a HER2-high-expressing human pancreatic cancer cell line. A: The contribution of Tmab to ADCC against Capan-1 was confirmed. The effector/target (E/T) ratio was set at 10:1, and 10 μg/ml of Tmab and isotype IgG were used. B: To confirm the appropriate concentration of Tmab, ADCC with the indicated dose of Tmab was measured. The E/T ratio was set at 40:1. C: To determine the necessity for the preadministration of Tmab to target cells, Capan-1 was incubated with 10 μg/ml of Tmab for the indicated times. Cells were harvested and ADCC was evaluated. The E/T ratio was set at 40:1.

### Trastuzumab-mediated ADCC against the HER2-low-expressing human pancreatic cancer cell line

Next, Tmab-mediated ADCC against the HER2-low-expressing human pancreatic cancer cell line was examined. As shown in Figure 3-A, although the Tmab-mediated ADCC against Panc-1 was elevated, the level was significantly weaker compared with that against Capan-1. The results in a representative individual cell are shown in Figure 3-B. In all investigated NK cells, ADCC against both target cells increased with the increase in NK cell number.

**Figure 3.**
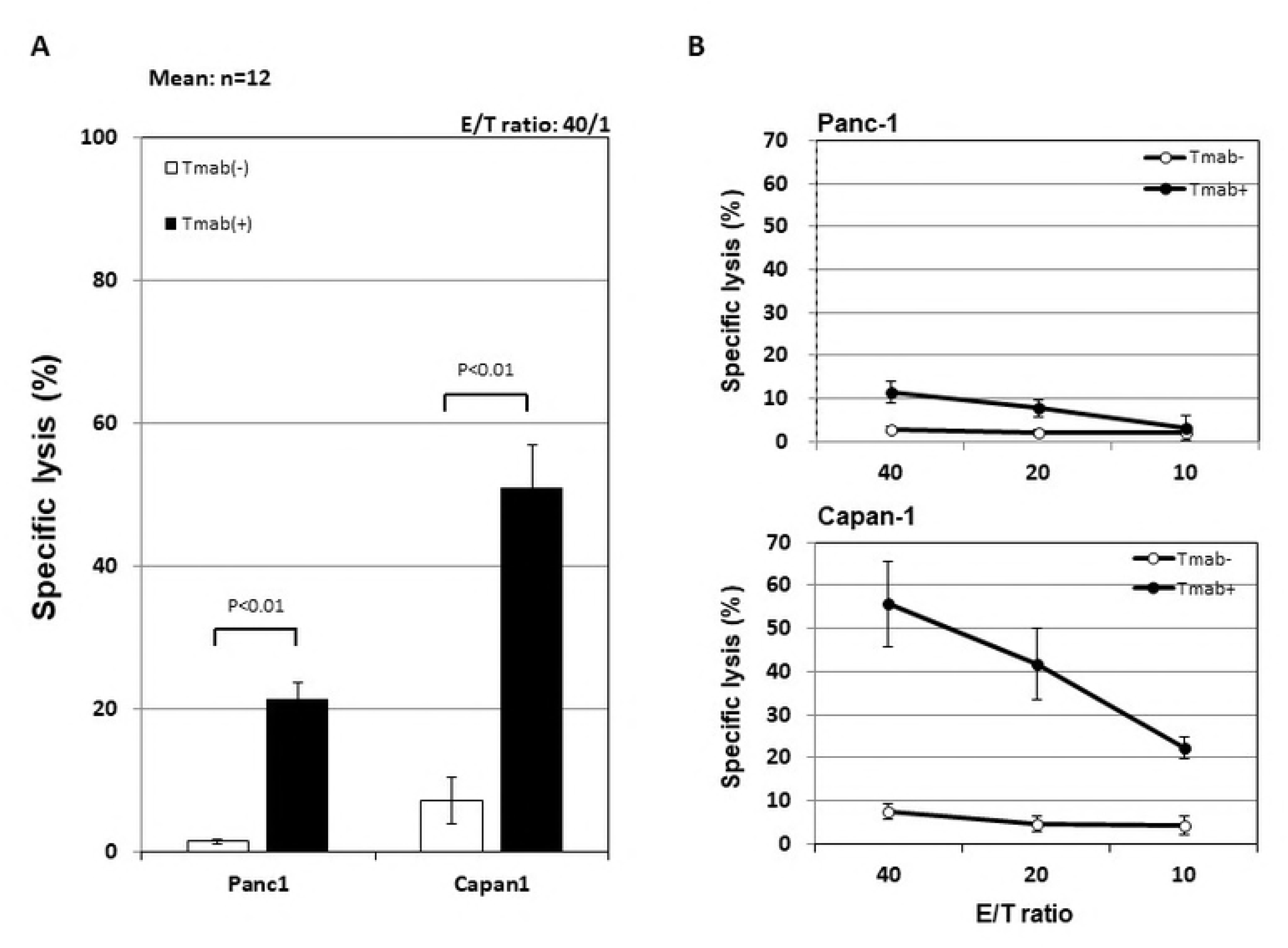
Tmab-mediated ADCC against Panc-1, a HER2-low-expressing human pancreatic cancer cell line, was investigated. A: Mean ADCC against Panc-1 was compared with that against Capan-1. B: ADCC against Panc-1 and Capan-1 in a representative individual was shown.

### Anti-CD137 mAb enhanced Tmab-mediated ADCC against the HER2-low-expressing human pancreatic cancer cell line

According to the results above, Tmab-mediated ADCC against Panc-1 could be improved. We therefore investigated the expression of various molecules, including CD137, associated with the activation of NK cells following Tmab treatment to determine which showed greater stimulation of NK cells. It was confirmed that only CD137 expression was up-regulated after Tmab stimulation (Figure 4-A). Moreover, the combination of αCD137 with Tmab enhanced Tmab-mediated ADCC against Panc-1 (Figure 4-B). However, the individual cell results showed that ADCC against Panc-1 was up-regulated when the expression of CD137 was increased following Tmab treatment (Figure 4-C), whereas it did not improve when CD137 remained unchanged (Figure 4-D).

**Figure 4.**
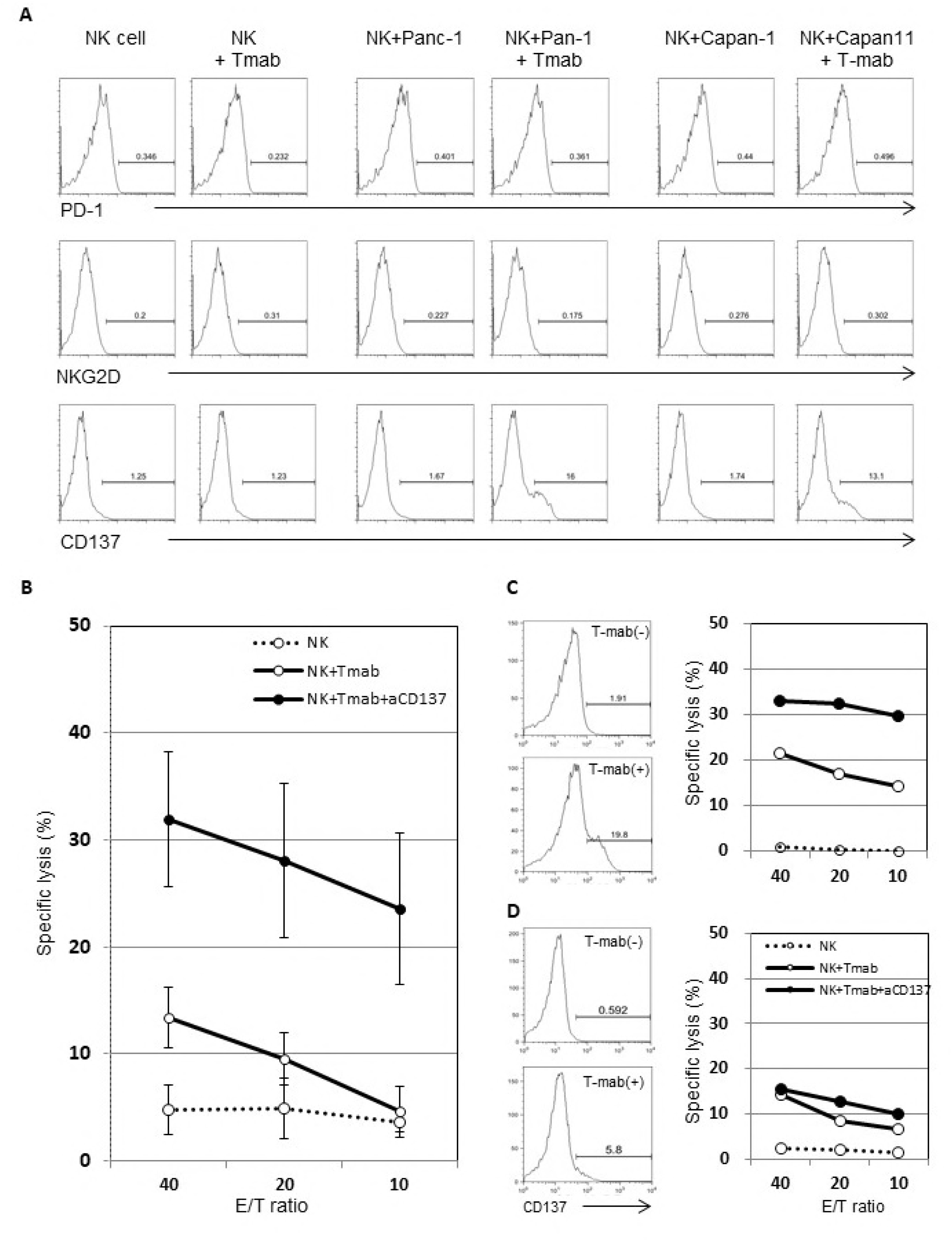
To establish how to increase ADCC against a HER2-low-expressing pancreatic cancer cell line, the addition of mAbs with Tmab was investigated. A: To select the optimal antibody, flow cytometric analysis was performed to evaluate changes in the various molecules on the surface of NK cells after Tmab administration. Based on the results, αCD137 was used in the following examinations. B: The effect of αCD137 combined with Tmab on ADCC against Panc-1 was evaluated. Mean ADCC in the indicated E/T ratios is shown. C: Results in a representative individual cell when ADCC increased with the addition of αCD137. Left two histograms: Changes in the expression of CD137 on the surface of NK cells. Right panel: Increase in Tmab-mediated ADCC with the addition of αCD137. D: Results in an unresponsive individual cell.

### Effects of the FcRγIIIA polymorphism on Tmab-induced ADCC against pancreatic cancer cell lines

It is known that the FcRγIIIA polymorphism is closely associated with the affinity to IgG-Fc [38], and the effects of the polymorphism against HER2-expressing breast cancer cells were reported [39]. FcRγIIIA 158V/V or V/F (VV/VF) conjugates easily with IgG-Fc to induce ADCC efficiently, whereas FcRγIIIA 158F/F (FF) conjugates weakly to IgG-Fc, resulting in weak ADCC. Based on those findings, we determined the FcRγIIIA polymorphism of each NK cell taken from 12 healthy individuals participating in this study and examined the effects on Tmab/αCD137-mediated ADCC against human pancreatic cancer cell lines. CD137 expression was more significantly elevated after stimulation with Tmab in both FcRγIIIA VV/VF individuals (n=8) than in the FF group (n=4) against both Panc-1 (Figure 5-A, left panel) and Capan-1 (Figure 5-A, right panel). Notably, the difference in CD137 up-regulation between the VV/VF and FF groups was striking when Panc-1 was treated with Tmab (Figure 5-A left panel). We additionally examined changes in IFN-γ (Figure 5-B) and TNF-α (Figure 5-C) released from NK cells in both the VV/VF and FF groups before and after treatment with Tmab in combination with αCD137 and found that the levels of these cytokines increased in the VV/VF group in the presence of Tmab. However, the addition of αCD137 did not affect the levels of these cytokines. The addition of αCD137 mAb significantly improved Tmab-mediated ADCC against Panc-1 (Figure 6-A, left panel) and improved more in VV/VF than in FF individuals, although the difference was not statistically significant (Figure 6-B, left panel). In contrast, although Tmab greatly enhanced ADCC against Capan-1 in both VV/VF and FF groups (Figure 6-A, right panel), the contribution of αCD137 was nonsignificant. Furthermore, the percentage of increase in ADCC did not differ between VV/VF and FF individuals (Figure 6-B, right panel).

**Figure 5.**
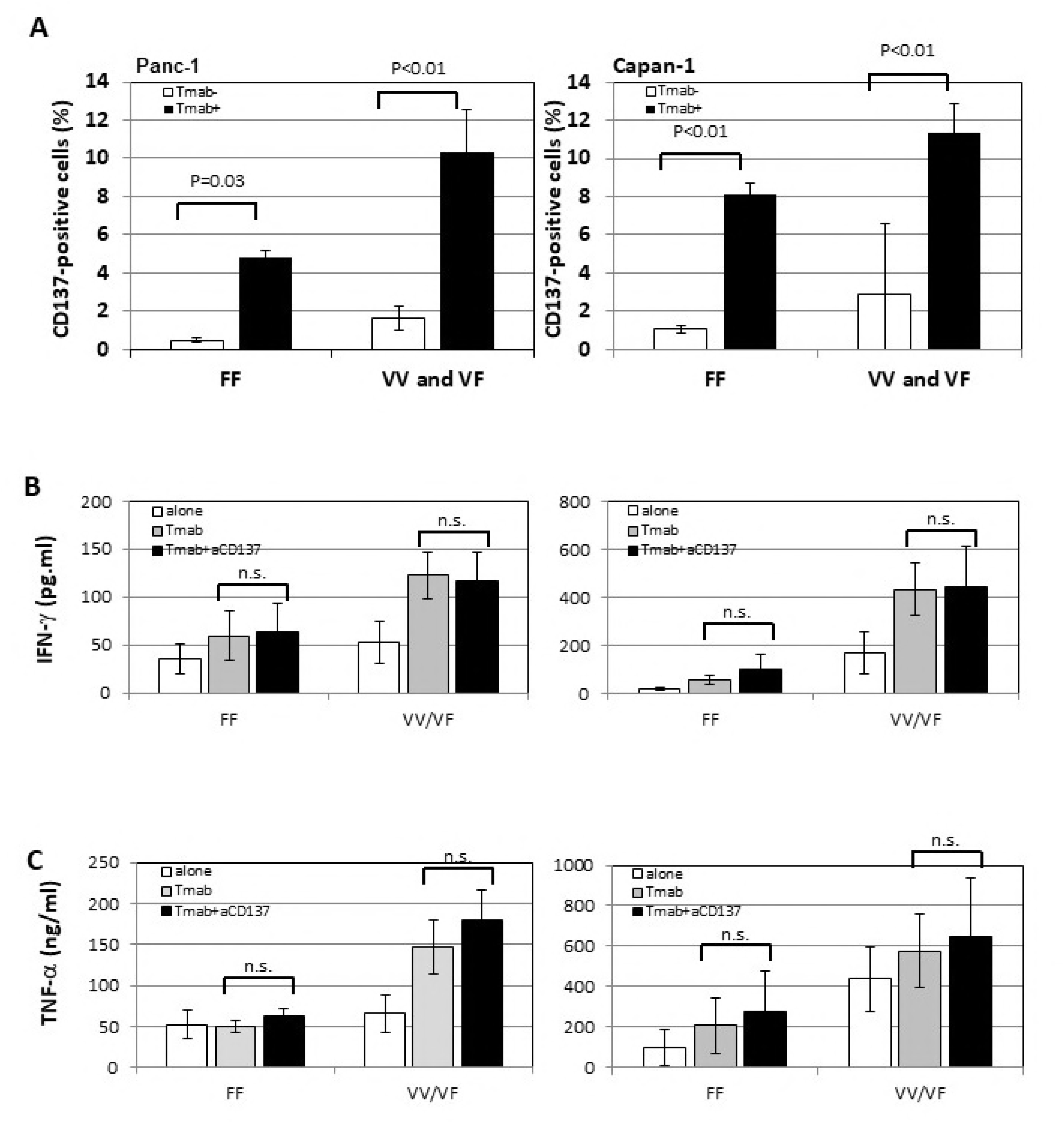
Effects of FcγRIIIA polymorphisms on the activity of NK cells. FcγRIIIA polymorphisms were determined by flow cytometric analysis and divided into the VV/VF (n=8) and FF groups (n=4). Differences in CD137 expression on the surface of NK cells in both groups were measured with flow cytometry at the end of Tmab-induced ADCC against Panc-1 (5A, left panel) and Capan-1 (5A, right panel). Levels of IFN-γ (5B, upper panels) and TNF-α (5B, lower panels) released from NK cells at the end of the ADCC assay were measured using ELISA.

**Figure 6.**
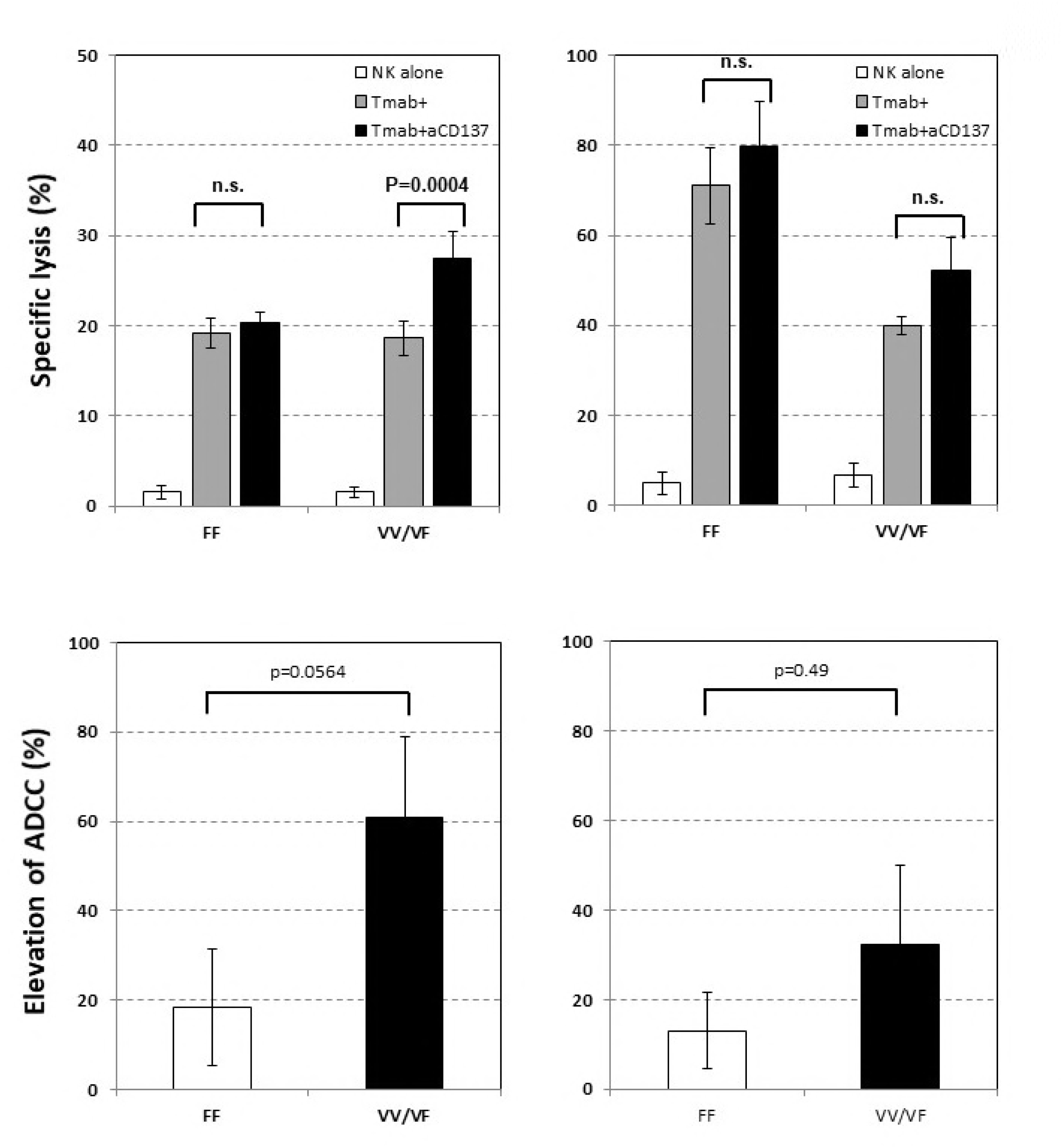
Effects of αCD137 addition to Tmab on the ADCC against Panc-1 (6A, left panel) and Capan-1 (6A, right panel) were compared between VV/VF and FF individuals. The percentage of increase in ADCC against Panc-1 (6B, left panel) and Capan-1 (6B, right panel) in each individual were calculated as indicated.

## Discussion

Our results indicated that Tmab can improve ADCC against a HER2-high-expressing human pancreatic cancer cell line. While Tmab-mediated ADCC against HER2-low-expressing human pancreatic cancer cell line was weak, it could be improved in the presence of αCD137 stimulating mAb to some degree. The FcRγIIIA polymorphism affected the activity of NK cells and was also associated with the level of Tmab-mediated ADCC against the HER2-low-expressing pancreatic cancer cell line. However, it did not exhibit optimal Tmab-mediated ADCC against the HER2-high-expressing pancreatic cancer cell line.

Tmab-mediated ADCC against the HER2-high-expressing cell line Capan-1 was potent without the addition of αCD137 mAb and independent of the FcγRIIIA polymorphism, indicating the clinical potential of Tmab for treating this type of pancreatic cancer. However, because HER2 expression on human pancreatic cancer cells is generally low, few patients are expected to benefit from a single administration of Tmab. Thus, additional techniques are needed to establish Tmab-related therapy for HER2-low-expressing cancer cells, the majority of human pancreatic cancer.

Our results indicated that the addition of αCD137 mAb enhanced Tmab-mediated ADCC against the HER2-low-expressing pancreatic cancer cell line. A previous report showed that Tmab-mediated ADCC against HER2-expressing human breast cancer cell lines was about 30% when they were exposed to a 20-fold amount of allogeneic NK cells [40]. Compared with that report, the cytotoxicity of Tmab plus αCD137 mAb-mediated ADCC against the HER2-low-expressing human pancreatic cancer cell line was slightly inferior to that against HER2-high-expressing human breast cancer cell lines because more NK cells were required to achieve the same level of ADCC in the former. Therefore increasing the number of NK cells in vivo will be crucial to establish combined therapy using Tmab and αCD137 mAb for the treatment of HER2-low-expressing human pancreatic cancer.

It was shown that intravenous administration of paclitaxel in breast cancer patients could recruit NK cells in the periphery [41]. The mechanism by which paclitaxel increases the number of peripheral NK cells is not known, although prior administration of paclitaxel is thought to up-regulate Tmab-mediated ADCC in the presence of αCD137 mAb against HER2-low-expressing human pancreatic cancer.

We also found that the binding affinity of FcRγlIIA to IgG-Fc is strongly associated with the expression of CD137 on the surface of NK cells along with its activation after stimulation with Tmab. Previous reports showed that the FcRγIIIA polymorphism is closely associated with the level of Tmab-mediated ADCC against human HER2-expressing breast cancer cell lines [38, 39]. Our results indicated that the FcRγIIIA polymorphism could also affect Tmab-mediated ADCC against the HER2-low-expressing human pancreatic cancer cell line, which seemed to be one of the reasons for the individual difference of ADCC levels against them. CD137 expression on the surface of NK cells with FcR-VV or -VF, which conjugates readily with IgG-Fc, was increased to enhance NK cell cytotoxicity in association with Tmab administration, resulting in improved ADCC against the HER2-low-expressing pancreatic cancer cell line. However, the prevalence of FcRγIIIA-VV was reported to be 7–10%, that of - VF 34–51 %, and that of -FF 42–56% in Japan [42, 43]. The prevalence of FcRγIIIA-VV/VF/FF in this study was 50.0/16.7/33.3%, however. Only a small cohort was analyzed in this study, which could explain the discrepancy, and more NK cells must be analyzed to clarify the relationship between FcRγIIIA polymorphisms and the ADCC against human HER2-low-expressing pancreatic cancer cells.

Based on our present and previous results, approximately 40–60% of pancreatic cancer patients can be expected to benefit from Tmab plus αCD137 mAb combined therapy. It remains unclear how the efficacy of the combination therapy could be improved in FcRγIIIA-FF individuals, who comprise the major population in Japan. Further investigations will be needed to resolve this. In addition, levels of IFN-γ and TNF-α released from NK cells after stimulation with Tmab increased more in VV/VF than in FF individuals, and there was no significant increase in the production of these cytokines after the addition of αCD137 mAb. The reason why activated, CD137-expressing NK cells did not secrete higher levels of cytokines after stimulation with αCD137 mAb was not determined in this study. More NK cells will be examined to resolve this.

For induction of effective ADCC by Tmab plus αCD137 mAb, a sufficient number of activated NK cells are necessary in patients with high-affinity polymorphisms against IgG-Fc. It will therefore be important to determine how to achieve this in vivo. The activity of NK cells is controlled by T-regulatory as well as T-helper lymphocytes [44], and gemcitabine, a standard reagent for treating unresectable pancreatic cancer, exhibits inhibitory activity against T-regulatory cells by down-regulating myeloid-derived suppressor cells that have a role in augmenting T-regulatory lymphocytes [45]. As described above, paclitaxel increases NK cells in the periphery immediately after administration, and they would be activated with the addition of gemcitabine. Thus, the newly established nab-paclitaxel regimen plus gemcitabine combination therapy appears to have the potential to increase activated NK cells in vivo, and the addition of Tmab and αCD137 is expected to improve the efficacy of that regimen against pancreatic cancer. Further investigations will be needed to confirm this hypothesis.

## Acknowledgment

We are grateful to Eiji Shinya, M.D., Ph.D. for his helpful suggestions on statistical analysis in this study.

